# Continental-scale integration of soil metagenomes and organic matter chemistry reveals ubiquitous microbial capacity for chemically-recalcitrant carbon decomposition

**DOI:** 10.1101/2025.07.03.663048

**Authors:** Young C. Song, Cheng Shi, Kelly G. Stratton, Christian Ayala-Ortiz, Izabel Stohel, Viviana Freire-Zapata, Malak M. Tfaily, Emiley Eloe-Fadrosh, Emily B. Graham

## Abstract

Soil organic matter (SOM) decomposition by microorganisms represents one of the largest uncertainties in predicting terrestrial carbon-atmosphere feedbacks. Yet we lack systematic understanding of the microbial taxa that drive decomposition of chemically distinct carbon pools and of the metabolic pathways they employ across environmental gradients. Here, we address this critical gap using the first continental-scale soil dataset pairing shotgun metagenomes with high-resolution molecular characterization of SOM chemistry. We use data from 47 standardized soil cores – selected using microbial respiration rates from 106 soils across the United States – to assemble 0.76 terabases (Tbs) of prokaryotic metagenome-assembled genomes (828 MAGs) and identify 66,727 distinct SOM molecules. Integrating these datasets revealed widespread microbial potential for depolymerizing chemically-recalcitrant carbon compounds previously considered stable. Notably, we uncover complementary metabolic specialization between genera affiliated with two abundant bacterial orders, *Rhizobiales* and *Chthoniobacterales,* and an archaeal order, *Nitrososphaerales.* This metabolic partitioning is consistent across soil depths and activity levels, suggesting coordinated decomposition of chemically-complex carbon through distinct but complementary biochemical strategies. The metabolic potential for depolymerization of chemically-recalcitrant compounds is supported by the abundance of these molecules across the soils, as indicated by Fourier-Transform Ion Cyclotron Resonance Mass Spectrometry (FTICR-MS), and by flux balance analysis of metabolic models. Our continental-scale integration of microbial genotypes with their chemical substrates reveals that a substantial fraction of SOM that is considered to be relatively stable harbors decomposition potential that current Earth system models fail to capture. These findings provide mechanistic insight for incorporating microbial metabolic pathways into carbon cycle predictions and highlight an underappreciated vulnerability of soil carbon to environmental variability.

## Main

Soil microbial respiration is a critical component of the global carbon cycle, yielding 98 ± 12 Petagram (Pg C/year) of carbon dioxide (CO_2_) into the atmosphere [1]. Most soil carbon (C) is contained within chemically diverse soil organic matter (SOM) pools that, through their interactions with plants, mineral surfaces, water, and other biological and physical soil properties, have the potential to sequester carbon within soils for centuries [2].

Chemically-recalcitrant SOM is typically thought to have long residence times in soils [3,4], and the widespread potential for its decomposition suggests a possible missing link in the global carbon cycle. Microbial metabolism of SOM proceeds through myriad metabolic processes [5], yet how interconnected factors and environmental contexts regulate SOM decomposition is largely unknown.

SOM contains a wide variety of organic molecules derived from plant-microbe interactions [5,6], biomass decomposition, and biochemical transformation into chemically-recalcitrant components [7,8]. These small ( <1,000 Dalton (Da)) compounds, including amino acids, lipids and organic phosphates, are transformed by soil microorganisms into CO_2_ through respiration [5]. While phylogenetic and metabolic inferences of soil metagenome-assembled genomes (MAGs) have revealed potential microbial mechanisms for transforming various organic molecules [9–11], robust characterization and confirmation of these pathways have historically depended on isolation and cultivation the organisms represented by MAGs [12]. Alternatively, organic matter characterization approaches using Fourier-Transform Ion Cyclotron Resonance Mass Spectrometry (FTICR-MS) provide molecular-level information about SOM chemistry [5], which can be integrated with MAG, potentially enhancing our understanding of SOM transformations while circumventing cultivation bottlenecks.

Given that the predicted residence times of stabilized SOM span decades to centuries, current Earth system models consider large fractions of SOM as carbon sinks [13]. We add to a growing body of work challenging this notion, while addressing knowledge gaps in the metabolic pathways that regulate decomposition of SOM across environmental gradients, using a unique soil dataset that integrates standardized metagenomics with high-resolution molecular characterization of SOM chemistry collected from surface and subsoils across the Continental United States (CONUS). Through the 1000 Soils Pilot of the Molecular Observation Network (MONet) [14], we assayed 106 surface and subsoils to uncover complementary roles in continental-scale soil C cycling processes conducted by genera within two predominant microbial clades, *Rhizobiales* and *Chthoniobacterales*. Our observations stress the vulnerability of the soil carbon stocks to increased microbial activity under continued environmental change, emphasizing the need for a tractable genetic framework for incorporating microbial mechanisms into Earth system models.

## Results

We investigated 106 soils that spanned the continental United States, representing 17 of 20 ecoclimatic domains defined by the National Ecological Observatory Network (NEON) [15]. This design enabled us to capture a wide range of spatial variation in soils across the continental United States and is consistent with other published continental-scale works [16,17]. We then assessed variation in microbial communities and SOM chemistry across 47 soils at two depths (surface and subsoil). Soils spanned 37 sites across the CONUS, encompassing most major ecoregions (Supplementary Fig. 1 and Supplementary Table 1). High-resolution mass spectrometry (FTICR-MS) identified 66,727 distinct SOM molecules (average 7.6 x 10^3^ molecules per sample), while metagenomic sequencing (average 1.1×10^8^ reads per sample) yielded 828 metagenome-assembled genomes (MAGs). Analyses of microbial relative abundance in soils indicated that on average 73.1% of metagenomic reads were microbial. Additionally, the assembled MAGs captured 50.6% and 36.5% of the bacterial and archaeal communities at the genus level, respectively.

While we acknowledge that non-prokaryotic organisms such as fungi and viruses have large impacts on biogeochemistry, assembling their metagenomes from soil remains a challenging task for several reasons, including short-read sequencing biases, limited reference databases, and variability in GC content [18–21]. These challenges were evident in our results, as taxonomic classification using Kraken2 [22] revealed that only a small fraction (less than 1%) of the reads from all soil samples were fungal. Therefore, we focused on bacterial and archaeal communities owing to their substantial roles in soil carbon cycling [23]. Given the above, computational approaches for soil microbiomes allow for deeper investigation into bacteria and archaea than other clades [18–21], and we expect that the results presented here will lay a foundation for investigations into the role of interkingdom interactions on belowground biogeochemistry, as new technological advances in identifying fungi and viruses are available. Detailed procedures of both SOM measurements and MAG assemblies are highlighted in the **Methods** section.

### Prevalence of Rhizobiales, Chthoniobacterales, and Nitrososphaerales across soil samples

We recovered 828 MAGs, which were dereplicated into 358 genomes. From the dereplicated MAGs, 48 were high quality (completion > 90% with contamination < 5%) and 310 were medium quality (completion ≥ 50% with contamination of < 10%) according to the Minimum Information about Metagenome-Assembled Genome (MIMAG; Supplementary Table 2) [24]. Bacterial MAGs constituted 89.1% of the dereplicated genomes (n=319 MAGs), while the remaining 39 MAGs were archaeal. Phylogenetic inference of the bacterial MAGs (143 from the surface and 176 from the subsoil) revealed that bacterial diversity was dominated by members of *Pseudomonadota* (21.0%; n=67), *Actinomycetota* (23.8%; n=76) and *Acidobacteriota* (17.2%; n=55). Of the archaeal MAGs, 92.3% (n=36) were classified as the phylum *Thermoproteota*, while the remaining three genomes were assigned to the phylum, *Thermoplasmatota*. Nearly 92% of the *Thermoplasmatota* genomes (n=34) were associated with the order *Nitrososphaerales* (Fig. 1).

**Figure 1.**
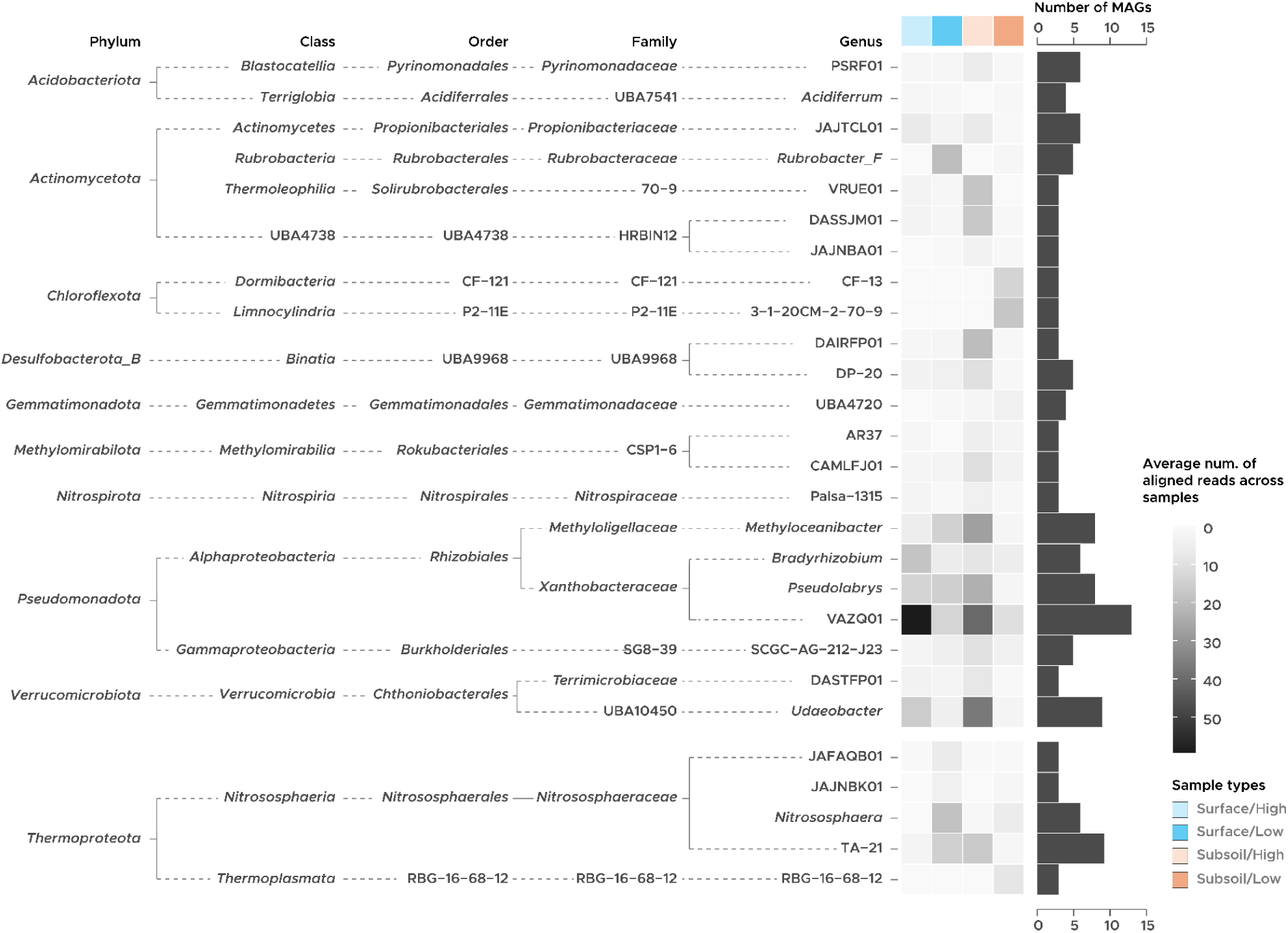
Distribution of bacterial and archaeal MAGs across soil layers and rates of potential microbial respiration. The bar chart shows the number of the dereplicated MAGs that was assigned to each genus-level lineage, while the intensity of the heatmap represents the average numbers of the reads from each soil types that were mapped to the genomes.

Examining soil microbial diversity at the order level revealed that *Rhizobiales* (phylum *Pseudomonadota*) and *Chthoniobacterales* (phylum *Verrucomicrobiota*) were the two most prevalent lineages, representing 12.0% (n = 43) and 6.1% (n = 22) of the dereplicated genomes, respectively (Fig. 1). Relative abundances inferred from read mapping indicated that some genera, such as the candidate group VAZQ01 (order *Rhizobiales*) and the *Udaeobacter* (order *Chthoniobacterales*) were the dominant taxa in high-respiration soils. Both VAZQ01 and *Udaeobacter* were also the two most prevalent genera overall, representing 3.6% (n=13) and 2.5% (n=9) of the total dereplicated MAGs, respectively. Genera within *Rhizobiales*, such as *Bradyrhizobium* and *Methyloaceanibacter*, were abundant across all soil types except for low-respiration subsoils.

Unlike *Rhizobiales* and *Chthoniobacterales*, some genera within the phyla *Actinomycetota* and *Chloroflexota* were more prevalent in the subsoils compared to the surface soils. For instance, the candidate genera VRUE01 and DASSJM01, both the members of *Actinomycetota*, had the highest average number of mapped reads in high-respiration subsoils, while the candidate groups CF-13 and 3−1−20CM−2−70−9 affiliated with *Chloroflexota* were the most prevalent genera in low-respiration subsoils. The only exception to this observation was the genus *Rubrobacter_F* within *Actinomycetota*, which was the most prevalent genus in low-respiration surface soils.

Of the genera within the archaeal order *Nitrosophaerales*, the candidate genus TA-21 and *Nitrosphaera* were the most prevalent groups, represented by 9 and 7 dereplicated MAGs, respectively. Both genera had the highest average number of mapped reads in low-respiration surface soils, while TA-21 was also present in high-respiration subsoils.

### Soil chemistry reveals depth-stratified carbon pools with distinct decomposition signatures

Pairwise comparisons of SOM molecular composition across depths and potential rates of microbial respiration revealed systematic chemical differentiation with implications for carbon cycling (G-test of uniqueness; Fig. 2 and Supplementary Table 3). While lignin- and condensed hydrocarbon-like compounds were prevalent across all soil types, we observed distinct SOM chemistry between surface and subsoils. For instance, comparison of chemistry in low-respiration subsoils with surface soils (both high- and low-respiration) revealed unique compounds in the subsoils (G-test p-value < 0.05; Fig. 2C and E), of which > 70% were composed of lignin-, condensed-hydrocarbon- and tannin-like compounds.

**Figure 2.**
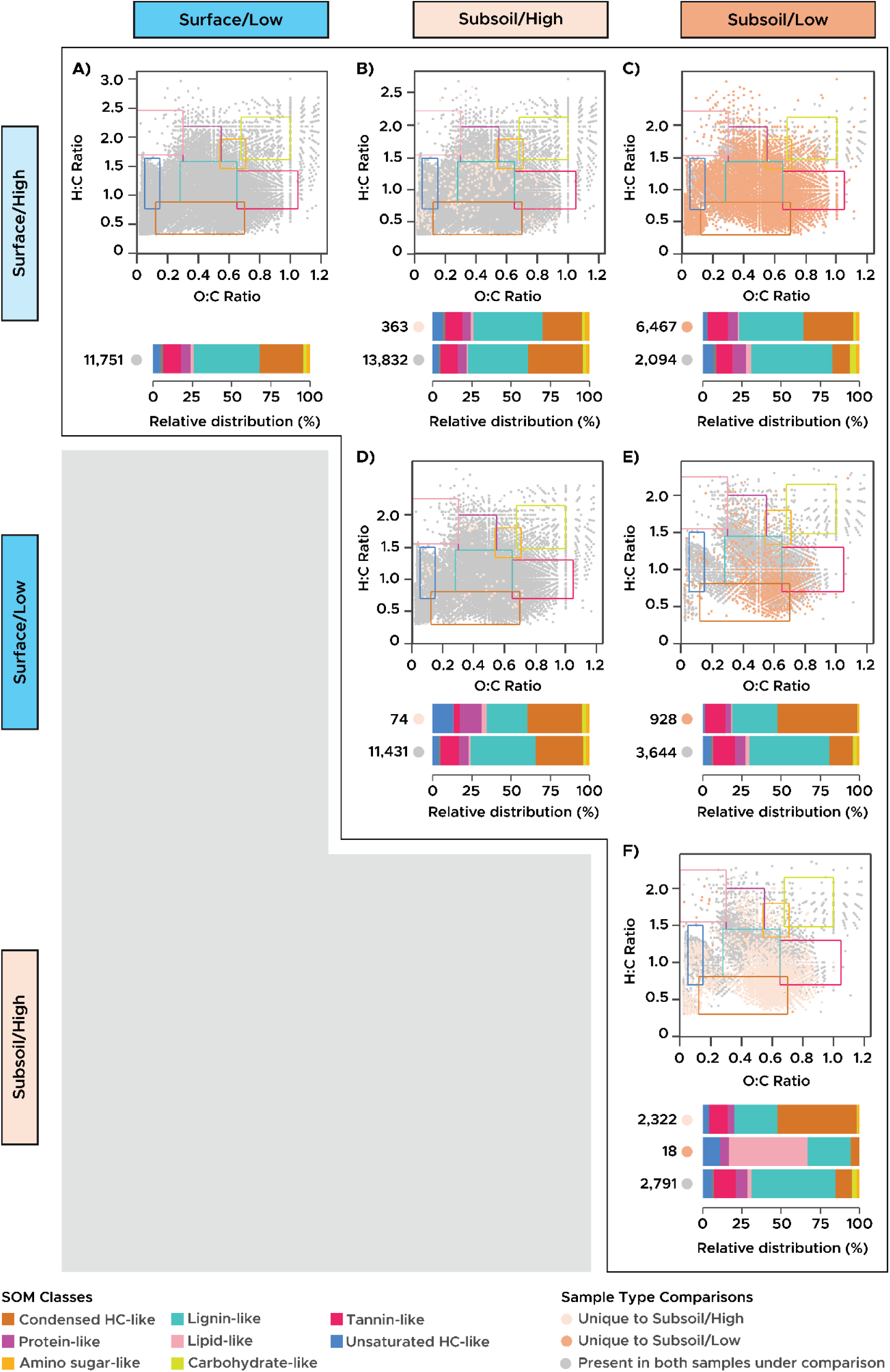
Molecular compositions of the SOMs across soil types according to the pairwise G-test of uniqueness. For each comparison, compounds unique to a soil type (p < 0.005) are represented as dots with colors other than gray, as specified in the legend. The boxes with colored edges on each van Krevelen plot denote the SOM classifications determined by thresholds of H:C and O:C ratios. The numbers located beside the stacked bar charts indicate the total number of compounds unique to each soil type being compared. The sizes of the colored segments in each bar represent the distribution (as percentages) of SOM classes among the total unique compounds in each soil type. The comparisons between the soil samples types are as follows: A) Surface/High vs. Surface/Low; B) Surface/High vs. Subsoil/High; C) Surface/High vs. Subsoil/Low; D) Surface/Low vs. Subsoil/High; E) Surface/Low vs. Subsoil/Low; and F) Subsoil/High vs. Subsoil/Low

Similarly, when comparing the chemistry of high-respiration subsoils against high-respiration surface soils, we observed SOM molecules unique to the subsoils, mostly composed (∼80%) of lignin-, condensed hydrocarbon-, and tannin-like compounds (Fig. 2B). Comparison between high-respiration subsoils and low-respiration surface soils (Fig. 2D) revealed that while 57% of SOM molecules unique to the subsoils were lignin-, condensed hydrocarbon- and tannin-like compounds, 32.4% were protein- (16.2%) and unsaturated hydrocarbon- (16.2%) like compounds (Fig. 2 and Supplementary Table 3). Between subsoil environments (Fig. 2F), high-respiration subsoils contained 2,322 unique compounds (>75% lignin- and condensed hydrocarbon-like molecules), while low-respiration subsoils had only 18 unique compounds, half of which were lipid-like—potentially reflecting microbial necromass under carbon-limited conditions.

### Microbial potential for depolymerizing chemically-recalcitrant SOMs is widespread

Using co-occurrence networks between annotated orthologs from MAGs and SOM molecules acquired from each of the four depth/respiration partitions, we evaluated potential relationships between microorganisms and metabolites. In particular, we identified densely connected modules within a large SOM-KEGG Orthology (KO) network (for all soil layers and potential respiration rates), revealing genes that showed significant relationships with chemically-recalcitrant classes of SOM, such as lignin-, condensed hydrocarbon-, and tannin-like molecules. A case in point is a set of 22 modules recovered in the high-respiration surface soils, with MCODE cluster scores ranging from 2.7 to 19.9 (Fig. 3 and Supplementary Fig. 2). We also recovered a single module from the low-respiration surface soil with a cluster score of 94.2. This module was composed of 189 nodes, 103 of which represented KO-annotated genes that were associated with metabolic pathways such as amino acid metabolism/synthesis, metabolism of C5 sugars, carbon fixation and cellular signalling and processing (Supplementary Table 4). Nearly 75% of the 86 SOM nodes within this module represented lignin-like compounds, while condensed-hydrocarbon- and tannin-like compounds were represented by 6 nodes each. This module also contained 5 amino-sugar-, 4 protein-like compounds and an unclassified compound.

**Figure 3.**
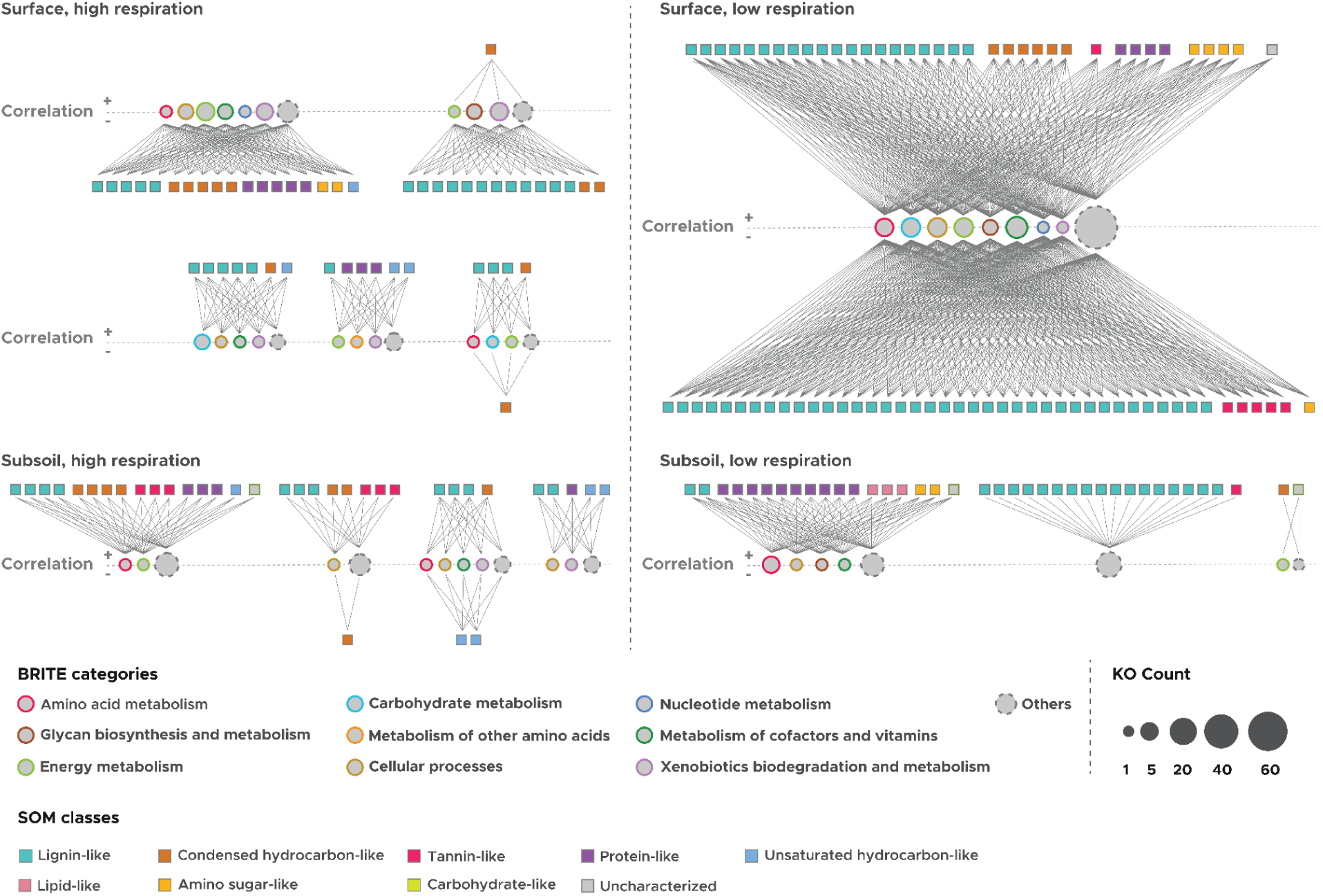
Modules with significant correlations between SOMs and KEGG Orthology (KO)-annotated genes across the soil layers and respiration rates, according to the MCODE network analysis. The shapes of the nodes within each module signify either KOs or SOMs, with their colors representing KEGG pathway categories for KOs or SOM classes, as detailed in the legend. Positive correlations between SOMs and KOs are depicted as edges situated above the dashed “Correlation” line, whereas edges indicating negative correlations are positioned below the dashed line. These correlations meet the criteria of rho > 0.6 and FDR-corrected P < 0.01.

The modules identified in high-respiration surface soils revealed the microbial potential for interacting with chemically-recalcitrant carbohydrates, including molecules resembling unsaturated hydrocarbons. Among the 22 modules detected in these soils, 15 contained between 2 and 5 KO-annotated genes linked to pathways such as amino acid catabolism, C5 sugar and starch metabolism, and secondary metabolite biosynthesis (Fig. 3 and Supplementary Fig. 2). Notable genes included *ligXa* and *graA* that are associated with the breakdown of lignin and other chemically-recalcitrant compounds [25,26]. These genes showed positive correlations with various combinations of metabolites, including one or multiple instances of lignin-, condensed hydrocarbon-, protein-, and unsaturated hydrocarbon-like compounds.

At the opposite end of our spectrum of soils, the module describing low-respiration subsoils was consistent with its unique chemistry profile, which included lipid-like compounds that were absent in the surface soils and in high-respiration subsoils (Fig. 3 and Supplementary Table 4). In this module, we observed 306 separate interactions (all represented by edges with correlations of > 0.95) between a set of SOMs including two lignin-like compounds, 10 protein-like compounds, three lipid-like compounds, two amino-sugar like compounds and an unclassified molecule, and 17 KO-annotated genes (Supplementary Table 4). Five of these genes were associated with amino acid catabolism, while the remaining 12 were linked to cellular functions (e.g., secretion system and biosynthesis of secondary metabolites and cofactors).

Genes in the KEGG database that are associated with metabolites detected by FTICR-MS (see **Methods**) further supported the potential for widespread microbial interactions with the chemically-recalcitrant SOM pool and its intermediates [27,28]. For instance, we uncovered metabolites associated with the biosynthesis of cofactors (map01240), which constituted 9.7%, 11.3% and 12.2% of metabolite-associated pathways identified in high- and low-respiration surface soils and in high-respiration subsoil, respectively (Fig. 4). We also detected metabolites affiliated with the metabolism of xenobiotics (map00980) and degradation of aromatic compounds (map01220) –– key pathways for processing complex organic polymers. These specialized metabolisms were distributed across all soil environments, constituting between 3.5% to 8.3% of metabolite-associated pathways in all groups of soils (Fig. 4).

**Figure 4.**
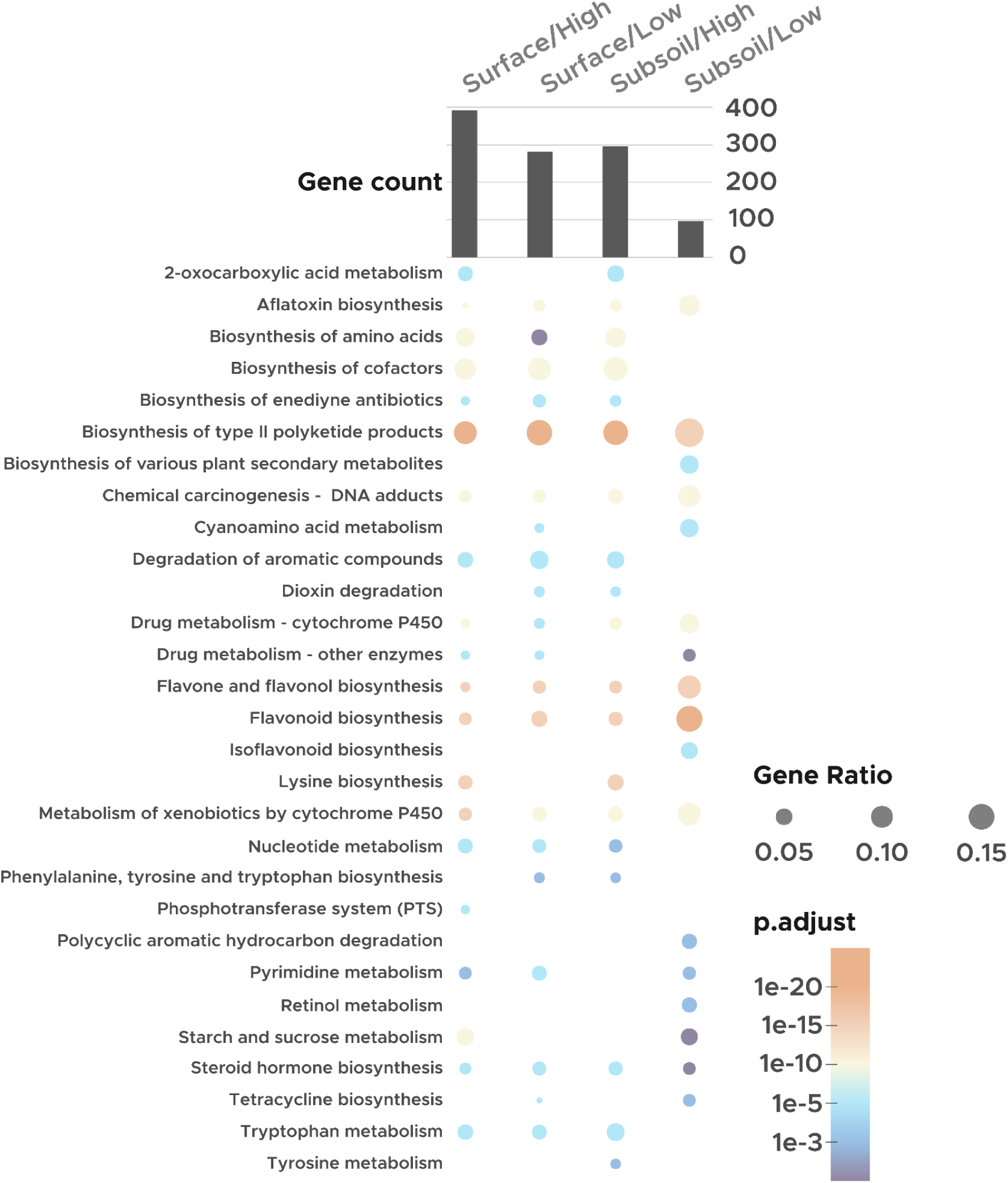
Distribution of high-level BRITE pathway categories based on recruitment of KO-annotated genes mapped to the FTICR-MS derived compounds. The histogram depicts the number of genes detected in the soil samples. The sizes of the bubbles indicate the relative distribution of genes associated with the designated BRITE pathway category, expressed as a proportion of the total gene counts in each sample. The adjusted p-values, represented by the colors of the bubbles, were calculated using the ClusterProfiler R package.

### Soil microbial communities universally encode complementary arsenals for depolymerizing chemically-recalcitrant SOM

Genomic analysis for carbohydrate-active enzymes (CAZymes) [29] revealed that the selected genera within the two widespread bacterial orders (*Rhizobiales* and *Chthoniobacterales*) and the archaeal order *Nitrososphaerales* possessed extensive and complementary enzymatic machinery for processing chemically-recalcitrant carbon substrates. Consistent with the prevalence of lignin-, tannin-, and/or hydrocarbon-associated SOM, these genera contained a variety of genes encoding complex carbon depolymerization (Fig. 5). For instance, we detected 92 CAZyme classes in ≥ 50% of the MAGs assigned to the archaeal genus, *Nitrososphaera*, while recovering 43, 60, 52 and 54 CAZyme classes in ≥ 50% of the *Methyloacenibacter*, *Bradyrhizobium*, *Pseudolabrys* and VAZQ0, respectively. In addition, 28 CAZyme classes were recovered in ≥ 50% of the genus *Udaeobacter* (Table 1). In all sets glycoside hydrolases (GH) and glycosyltransferases (GT) were the most prevalent CAZyme classes. Unlike the bacterial genera, *Nitrosphaera* and the *Candidatus* TA-21 also possessed polysaccharide lyases (PL).

**Figure 5.**
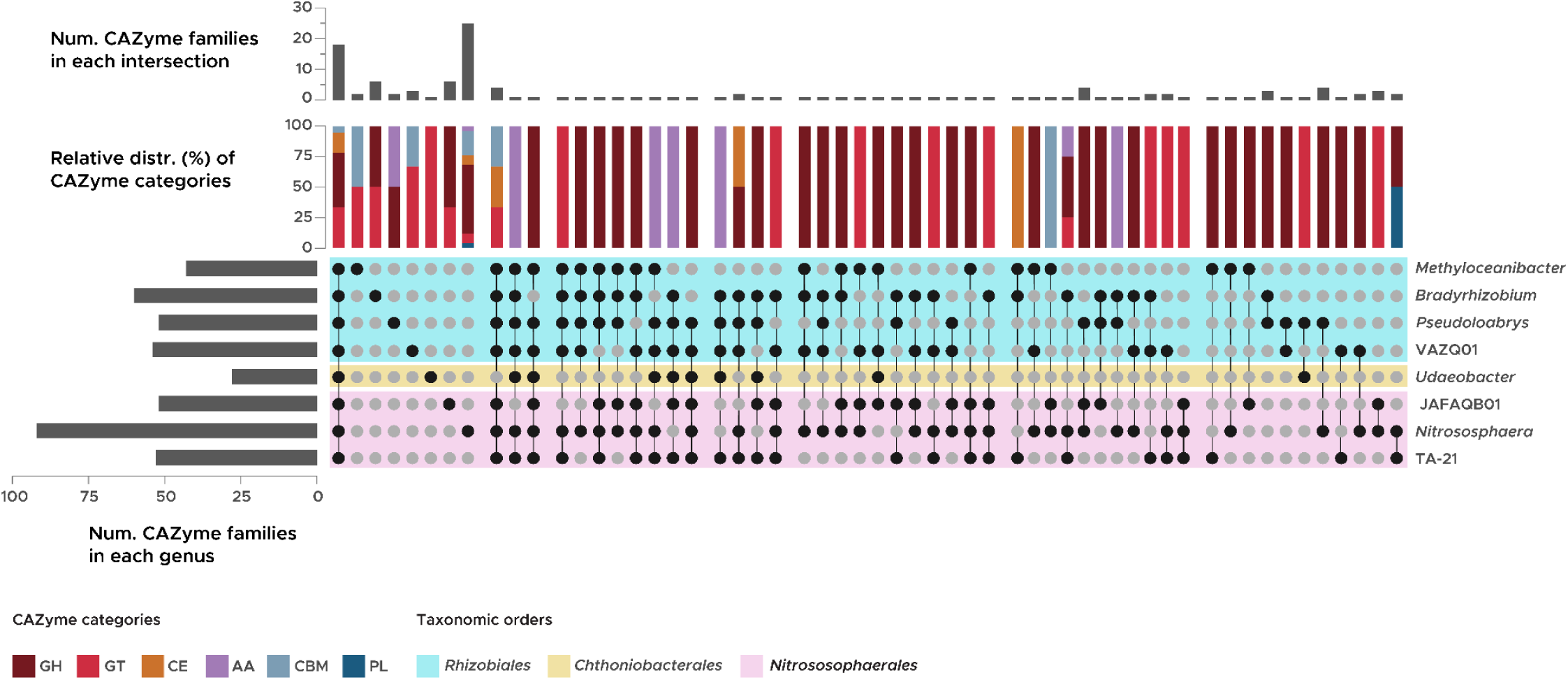
Carbohydrate-active enzymes (CAZymes) detected in MAGs of the selected genus-level lineages affiliated with the orders *Rhizobiales*, *Chthoniobacterales* and *Nitrososphaerales*. The bar chart on the left illustrates the total number of CAZyme families identified across the selected genera. Meanwhile, the combined dot-and-line plot along with the bar chart at the top displays the count of CAZyme families found in the specified genera or genus. Additionally, the stacked bar chart highlights the proportional distribution of CAZyme categories among these families. The CAZyme categories, as defined in the legend, include GH (glycoside hydrolase), GT (glycosyl transferase), CE (carbohydrate esterase), AA (auxiliary activity), CBM (carbohydrate-binding module), and PL (polysaccharide lyase).

A cross-comparison of the CAZymes identified in the MAGs from each selected genus further highlighted *Nitrosphaera*’s metabolic potential for degrading a wide variety of chemically-recalcitrant carbohydrates. Of the 92 CAZyme classes recovered from the genomes belonging to this genus, 25 classes were unique to *Nitrosphaera*. Of them, 52% (13) were GHs, with the rest classified as GTs, auxiliary activity enzymes (AAs), carbohydrate esterases (CEs), carbohydrate binding molecules (CBMs) or PL. The composition of the CAZyme classes in the other selected bacterial and archaeal genera were not as diverse as *Nitrososphaera*. For instance, of the 6 enzymes unique to the candidate archaeal genus JAFAQB01, 66.7% (4) were GHs, while the remaining two were GTs. Similar observations were made in the genera belonging to both *Rhizobiales* and *Chthoniobacterales*, where the majority of the CAZymes unique to each of these lineages were dominated by GHs, GTs and CBMs.

While the recovery of fungal MAGs from metagenomic sequencing was limited [18,21], we uncovered a set of potential fungal pathways for depolymerizing SOM from annotated contigs. In total, we detected 11 CAZymes that were present in fungal contigs in ≥ 50% of the soils, largely classified as GH, GT, AA or CBMs (Supplementary Fig. 3). Of these CAZymes, there was a single AA unique to the high-respiration surface soils and two GTs unique to the low-respiration subsoils. We also detected a GH unique to subsoils across both high- and low-respiration.

### Complementary amino acid and inorganic nitrogen acquisition mechanisms are enriched in *Rhizobiales* and *Nitrososphaerales*-associated genera

Flux balance analysis (FBA) of the selected genera belonging to *Rhizobiales*, *Chthoniobacterales* and *Nitrososphaerales* revealed that despite the observed metabolic potentials for amino acid catabolism in all prokaryotic MAGs, substrate preferences for these amino acids appeared to differ by genera (Fig. 6). A closer examination of the selected bacterial and archaeal MAGs revealed a set of genes encoding the transformation of L-aspartate, L-glutamate, L-arginine and L-serine to various intermediates of the TCA cycle (Supplementary Table 5). For instance, *Nitrosphaera* and *Udaeobacter* MAGs possessed at least a single copy of most genes encoding transformation of L-aspartate to L-asparagine or oxaloacetate. Meanwhile, the presence of the glutamine synthetase gene (*glnA*), which facilitates the amidation of L-glutamate to synthesize L-glutamine, was more prevalent in *Bradyrhizobium* compared to other genera. FBA results were supported by the distribution of KO-annotated genes, which showed metabolic potential for transforming both L-aspartate and L-glutamate in *Methyloaceanibacter* and *Pseudolabrys*. The annotation profiles for the bacterial and archaeal genera aligned with the FBA, which indicated a high average uptake rate of L-aspartate and/or L-glutamate for for *Nitrosphaera*, *Udaeobacter* and most genera in *Rhizobiales*.

**Figure 6.**
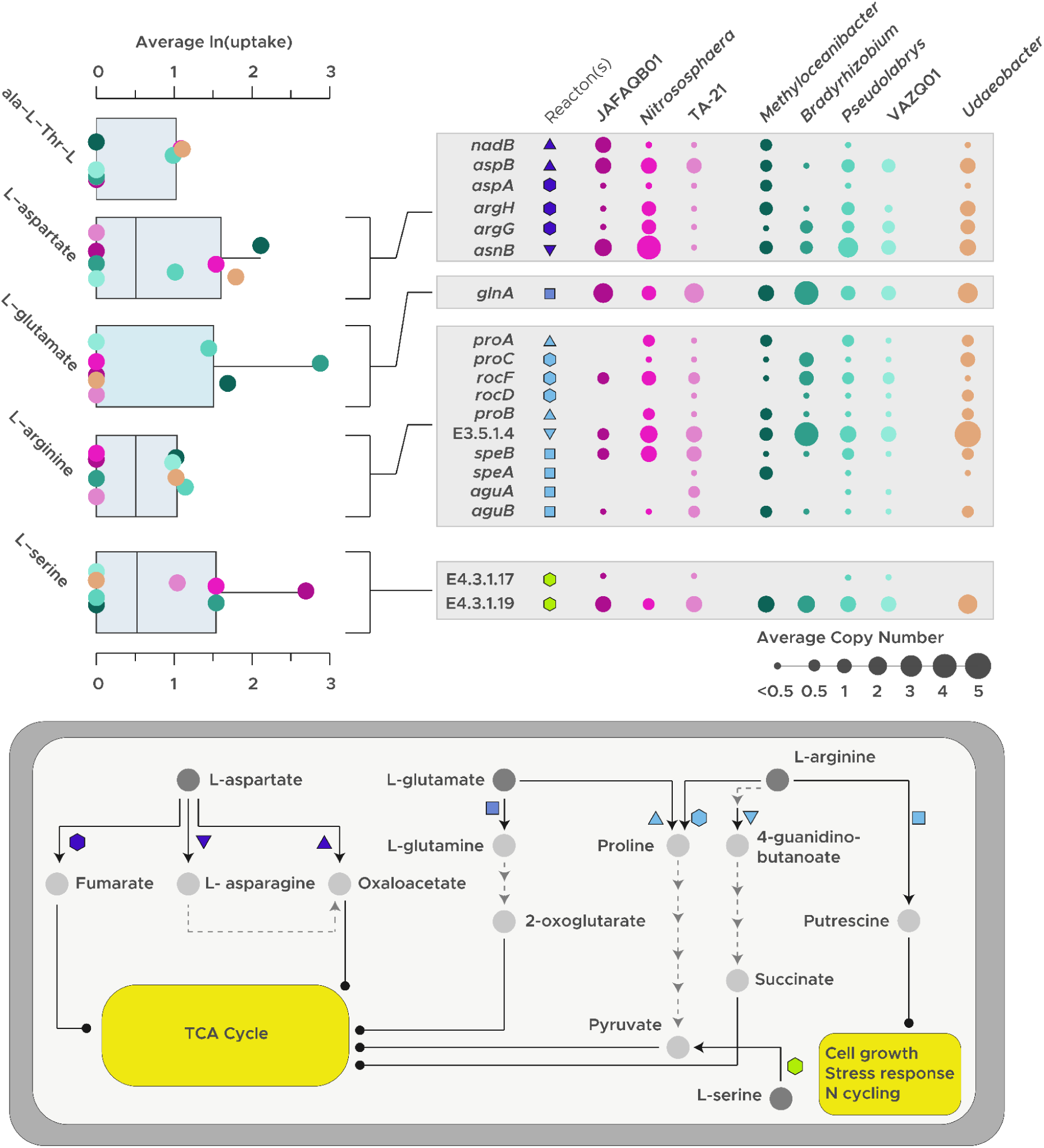
Metabolic potentials of amino acid catabolism in the selected bacterial and archaeal genera, based on the flux-balance-analyses (FBA) and the KO annotation of the protein coding genes detected in MAGs. The box plot in the top-left shows the average log-corrected uptake rates of amino acids for each genus, as determined through FBA, while the bubble plot in the top-right highlights the average copy numbers of each KO associated with the corresponding genes. The color-coding of the dots/bubbles in both plots represents one of the chosen bacterial or archaeal genera. In addition, the shapes within the bubble plot in the leftmost column denote specific reactions, which are depicted in the cell diagram at the bottom.

The transformation of L-arginine was notably prevalent among *Methyloaceanibacter*, *Pseudolabris*, candidate genus VAZQ01, and *Udaeobacter*, with these four bacterial genera exhibiting a higher average uptake of arginine compared to other lineages under study (Fig. 6). Interestingly, MAGs associated with *Bradyrhizobium* and *Udaeobacter* displayed a high average copy number of the gene encoding aliphatic amidase (*amiE*), an enzyme that catalyzes the hydrolysis of short-chain amides generated from the activity of arginine monooxygenase [30]. However, the average arginine uptake in *Bradyrhizobium* was close to zero, unlike in *Udaeobacter*. Additionally, most *Bradyrhizobium* MAGs lacked genes associated with the conversion of arginine to putrescine (e.g., *speB*, *speA*, *aguA*, and *aguB*), distinguishing them from other *Rhizobiales*-associated genera. Similarly, the same genes were absent in JAFAQB01 and *Nitrososphaera*. These archaeal genera, along with TA-21, exhibited an almost negligible average arginine uptake rate. All bacterial and archaeal MAGs contained at least one copy of *ilvA*, which encodes L-serine/L-threonine dehydratase, an enzyme responsible for converting serine into pyruvate. Among these genera, only the three archaeal genera and *Bradyrhizobium* demonstrated a comparable average serine uptake rate. These patterns suggest complementary amino acid utilization strategies between bacterial and archaeal lineages, whereby arginine catabolism is concentrated in genera such as *Methyloaceanibacter*, *Pseudolabrys*, candidate genus VAZQ01, and *Udaeobacter*, while archaeal taxa and *Bradyrhizobium* rely more consistently on serine, aspartate and glutamate rather than arginine-based pathways.

In addition to complementary amino acid catabolism, bacterial and archaeal lineages also displayed partitioned pathways for inorganic nitrogen transformations, with *Nitrosphaerales* specializing in ammonia oxidation and key bacterial genera encoding dissimilatory nitrate reduction. A nearly complete set of genes involved in ammonia oxidation to nitric oxide (NO), including *pmoABC* and *nirK*, was found across all three *Nitrosphaerales* genera (Supplementary Table 5). However, the *hao* gene, which encodes hydroxylamine dehydrogenase, was notably absent in all *Nitrosphaerales* MAGs. In contrast, ammonia oxidation genes were entirely absent from *Rhizobiales* and *Chthoniobacterales* MAGs. Instead, a partial set of genes involved in dissimilatory nitrate reduction, such as *narGHI*, *napAB*, *nirBD*, and *nrfAH*, were detected, particularly in *Methyloaceanibacter* and *Udaeobacter* (Supplementary Table 5).

### Lipid metabolism reveals contrasting survival strategies under carbon limitation

The distribution of SOM molecular classes across soils revealed that while microbial potential for depolymerizing chemically-recalcitrant organic matter appeared to be widespread in surface soils and in high-respiration subsoils, biomass synthesis pathways tended to dominate low-respiration subsoils. A complete or near-complete set of lipopolysaccharide biosynthesis genes (e.g., *lpxD*, *lpxL*, and *gmhC*) were identified in MAGs across nearly all bacterial and archaeal genera studied, with the exception of *Bradyrhizobium* (Supplementary Table 6). Notably, these genes were present in ≥ 50% of the MAGs affiliated with *Nitrososphaera* and *Udaeobacter*, highlighting fundamentally different strategies in these microbial lineages for surviving carbon limitation [31,32].

We also investigated the occurrence of lipid biosynthesis genes in MAGs associated with two candidate genera, CF-13 and 3-1-20CM-2-70-9, both classified under the phylum *Chloroflexota* and the most prevalent taxa in low-respiration subsoils (Fig. 1). Compared to MAGs of dominant bacterial and archaeal lineages, the genomes of CF-13 and 3-1-20CM-2-70-9 exhibited lower CheckM completion rates (between 50% and 80%). Only a partial set of lipopolysaccharide synthesis genes was detected in a small fraction of these MAGs, consistent with their relatively lower genome completion rates relative to those of more prevalent bacterial and archaeal lineages.

## Discussion

Our study provides evidence that soil microbiomes across the continental United States possess genetic potential associated with the depolymerization of compounds traditionally considered to be chemically stable, including complementary metabolic specialization among abundant bacterial (*Rhizobiales*, *Chthoniobacterales*) and archaeal (*Nitrososphaerales*) lineages. This widespread microbial capacity for chemically-recalcitrant SOM turnover underscores the need for further investigations into the specific enzymatic pathways (e.g., CT-depolymerization, C_15_ metabolism and phenolic metabolism [33]) that appear to support the breakdown of chemically-recalcitrant compounds in most soils. By analyzing 828 MAGs - dereplicated to 358 -dereplicated MAGs and 66,727 molecular-level SOM measurements, we identified genera within dominant bacterial and archaeal orders—*Rhizobiales*,*Chthoniobacterales* and *Nitrosospaherales*—that use complementary enzymatic strategies to process chemically complex carbon across diverse soils. These findings suggest that slow carbon pools in current models may be more vulnerable to decomposition than previously thought, exposing a critical gap in carbon-climate feedback predictions.

It is important to note that SOM degradation not only depends on the chemical composition of SOM, but also on factors that control how easily microbes can encounter and use that material [16,17,34,35]. The rate at which SOM turns over is governed both by the chemical properties of carbon inputs and by physical and spatial constraints that limit contact between microorganisms and organic substrates. Organic compounds that are chemically bioavailable could be protected from microbial decomposition by physical barriers that restrict access or create micro-environmental constraints on decomposer activity and movement—for example, protection within soil aggregates or nanopores, or isolation in ice or anaerobic microsites [36,37]. These abiotic factors themselves could also be influenced by chemistry (e.g., chemical binding). Together, these approaches emphasize that accurate representation of soil carbon dynamics will require coupling molecular-level descriptions of SOM chemistry with explicit consideration of microbial physiology and the spatially heterogeneous soil environment. Although we focus on dissolved SOM, particulate organic carbon also plays key roles in nutrient cycling, carbon storage, and the formation of stable organic matter fractions that underpin soil health and ecosystem sustainability [38,39]. Together, these perspectives emphasize that accurate representation of soil carbon dynamics will require coupling molecular-level descriptions of SOM chemistry with explicit consideration of microbial physiology and the spatially heterogeneous soil environment.

Integrating chemical profiling of SOM and functional annotation of soil MAGs showcases widespread microbial potential for lignin-, condensed hydrocarbon-, and tannin-like molecules that are typically thought to have long residence times in soils (Figs. 3 and 4) [40]. These dynamics may arise from microbial interactions with plant residues and their decomposition products, which are shaped by the molecular composition and spatial arrangement of SOM, as well as the functional potential of the co-located microbiome [40–42]. The prevalence of lignin-and condensed hydrocarbon-like molecules across soil depths in particular, regardless of microbial activity level, and their association with genes encoding SOM-degrading enzymes suggest current soil carbon stocks may be vulnerable to microbial activity in response to changing environmental conditions (Figs. 2, 3, and 4) [43,44]. The apparent persistence of the soil carbon pool therefore likely reflects limits on microbial access or activity, and the shifts in factors such as temperature, moisture, or redox conditions could relax these constraints and stimulate additional decomposition of existing soil carbon stocks. Identifying genes, such as *ligXa* and *graA*, directly linked to chemically-recalcitrant carbon processing highlights potential microbial mechanisms in decomposing chemically-recalcitrant carbohydrates, such as lignin to simpler molecules [25,26,45]. Additionally, genes for polyketide synthesis and other secondary metabolites may contribute to SOM depolymerization through mechanisms like metal mobilization [46,47] and the activation of quorum sensing activation [48]. The detection of complex networks in the low-respiration surface soils, together with prevalence of *Methyloceanibacter*, *Pseudolabrys*, and *Nitrososphaera* at the same soil depth and respiration level further underscores the increased reliance of these microorganisms on accessible carbohydrates during periods of microbial low activity (Figs. 1 and 3). While these bacterial and archaeal lineages are primarily known for their C1 and nitrogen metabolism (e.g., nitrogen fixation or nitrate reduction), presence of CAZymes for depolymerizing complex carbon polymers in some members of these clades have also been previously described [49–53]. Together, this suggests that under low microbial activity, communities in surface soils are supported by taxa such as *Methyloceanibacter* and *Pseudolabrys*, which encode diverse CAZymes and central carbohydrate pathways alongside aspartate-, glutamate-, and arginine-catabolic routes, and by *Nitrososphaera* with its capacity for ammonia oxidation, enabling continued turnover of accessible complex SOM even when bulk respiration is low.

The widespread presence of *Rhizobiales*, *Chthoniobacterales*, and *Nitrososphaerales* across all soils, combined with the abundance of CAZymes in the MAGs associated with genera within these orders, suggests mechanisms for the depolymerization of SOM during both high and low microbial activity in surface and subsurface soils (Fig. 5). Notably, the prevalence of GH and GT highlights their ability to interact with glycans, monosaccharides, and polysaccharides, in particular, and to construct extracellular structures that support adaptation to varying levels of carbon availability [54,55]. Additionally, the detection of carbohydrate-binding modules (CBMs) unique to genus-level lineages within *Rhizobiales*, such as *Methyloceanibacter* and VAZQ01, further emphasizes the potential of these microbes to break down chemically-recalcitrant compounds, particularly under low microbial activities.

*Nitrosphaera* in particular contained a distinct set of CAZymes, exhibiting greater diversity compared to those found in other bacterial or archaeal MAGs (Fig. 5). While members of this genus have traditionally been identified as ammonia-oxidizing chemolithotrophs, recent studies indicate metabolic versatility within certain *Nitrosphaera* sub-clades [49]. In support of this emerging idea, we detected both intracellular and extracellular CAZymes, such as CBMs and GHs, alongside genes involved in ammonia oxidation (Table 1 and Supplementary Table 5). Additional evidence for metabolic diversity in *Nitrosophaera* includes the presence of amino acid catabolic pathways, particularly for L-aspartate and L-serine (Fig. 6). Given the prevalence of *Nitrosophaera* in low-respiration surface soils, as well as in subsoils with limited respiration, we hypothesize that these archaea are equipped with mechanisms utilize amino acids as a source of nitrogen required to depolymerize complex carbon compounds, enabling them to persist under energy- and carbon-limited conditions.

In *Nitrosphaera* and other genera within *Nitrosphaerales*, pathways involving amino acids appear to be coupled with ammonia oxidation, given that none of the MAGs belonging to these groups possessed the *hao* gene, which is thought to be an essential component of ammonia oxidation [56–58]. The absence of *hao* may reflect sequencing limitations or suggest the involvement of uncharacterized Cu-protein complexes in ammonia oxidation, a mechanism observed in other ammonia-oxidizing archaea (Supplementary Table 9) [57]. Ammonia that is oxidized by *Nitrososphaerales* could be generated from catabolism of available amino acids in soil. These findings underscore the role of *Nitrososphaerales* in linking soil carbon and nitrogen cycles and highlight the need for further investigation into their genetic machinery for ammonia oxidation.

Amino acid and inorganic nitrogen acquisition mechanisms were also enriched in *Rhizobiales* and *Chthoniobacterales*, as observed in flux balance analyses of the MAGs, indicating a vital role of these pathways in nitrogen acquisition from SOM and energy generation in *Rhizobiales* and *Chthoniobacterales* in particular (Fig. 6, Supplementary Fig. 2) [59,60]. In *Rhizobiales*- and *Chthoniobacterales*-affiliated genera, particularly in *Methyloceanibacter* and *Udaeobacter*, intermediates generated through amino acid catabolism (e.g., pyruvate, NADH, and FADH2) could serve as sources of reducing power to drive dissimilatory nitrate reduction, which was also detected in these lineages (Supplementary Table 5). Therefore, the degradation of amino acids like L-arginine, observed in *Methyloceanibacter* and *Udaeobacter*, along with their prevalence in soils with low microbial activity, indicates that amino acid catabolism is crucial for maintaining energy balance and supporting nitrogen biosynthesis required for fundamental cellular functions (Figs.1 and 6) [59,61–63].

The broad spectrum of CAZyme classes identified in *Rhizobiales*-affiliated genera versus other MAGs highlights a putative advantage for them to break down chemically-recalcitrant SOM in soils with high microbial activity. This is especially evident for *Bradyrhizobium*, *Pseudolabrys*, and VAZQ01, which were prevalent in both surface and subsoils with elevated respiration rates (Figs. 1 and 5). While the potential to decompose complex carbohydrates has been documented for both *Bradyrhizobium* and *Pseudolabrys*, the metabolic characteristics of VAZQ01 remain poorly understood [64,65]. Although the *Chthoniobacterales*-affiliated lineage *Udaeobacter* was also present in both surface and subsoils with high respiration, the limited set of CAZymes detected in its MAGs indicates a dependence on mono- and polysaccharides for carbon and energy metabolism, further reinforcing its auxotrophy for various amino acids. (Figs. 5 and 6; [66]). The amino acid catabolism observed in *Udaeobacter*, combined with its previously identified ability to sustain the respiratory chain using atmospheric H_2_ under nutrient-deficient conditions, aligns with its prevalence in high-respiration subsoils, where carbon and nitrogen sources are scarce and competition for these resources is intense (Fig. 1; [67,68]). These findings suggest contrasting carbon usage strategies in *Rhizobiales* and *Chthoniobacterales*, with *Rhizobiales* utilizing a broader range of carbohydrate polymers and amino acids as sources of both carbon and nitrogen, whereas *Chthoniobacterales* relies on scavenging amino acids, likely derived as byproducts of microbial activities in soil, highlighting how taxon-specific metabolic traits can shape nutrient cycling and microbial interactions in terrestrial ecosystems.

Lastly, the bioavailability of soil lipids has been hotly debated, because they may dually serve as a high-energy substrate and as a reservoir of sequestered carbon derived from microbial necromass [69–71]. We found a diversity of lipid-like molecules in low-respiration subsoils that may represent sequestered plant and/or microbial residues from prior periods of higher metabolic activity [72,73] (Fig. 2). In subsoils, nutrient limitations and/or mineral protection likely promote the disproportionate accumulation of lipid-like molecules, contributing significantly to the stable fraction of SOM. While lipid persistence has been linked to soil carbon stabilization, the universality of soil microbial potential for lipid metabolism –– even in soils with low rates of measured metabolic activity –– suggest that they may become susceptible to decomposition as environmental conditions change. Notably, interactions between lipid-like molecules and microbial genes in low-respiration soils also indicate that lipid-like necromass components may be vulnerable to microbial decomposition during periods of reduced carbon availability and respiration rates [73,74] (Fig. 3). Thus, we highlight potential lipid turnover as a key consideration in soil carbon dynamics [51].

Strong network correlations between lipid-like compounds and microbial secretion systems, cofactors, and secondary metabolites in low-respiration subsoils may also elucidate strategies for microbial adaptation to carbon-limited environments (Figs. 3 and 4; Supplementary Table 6). Secretion systems enable microbial interactions by exporting compounds to facilitate nutrient acquisition, particularly labile substrates [75,76], while cofactors (e.g., molybdopterin) and secondary metabolites support energy metabolism, redox reactions, and oxidative damage mitigation under stress [77,78]. The functional annotations of the MAGs linked to *Rhizobiales*, *Chthoniobacterales*, and *Nitrososphaerales* suggest that these microbes may possess the ability to enhance lipopolysaccharide (LPS) synthesis, thereby stabilizing their cell envelope and boosting their resistance to environmental stress [79,80]. These characteristics were notably prevalent in *Udaeobacter* and *Nitrososphaera* (**Supplementary Table 6**). Gram-negative bacteria, including *Udaeobacter* and *Nitrososphaera*, contribute significantly to organic carbon storage through the chemical persistence of LPS, particularly its chemically-recalcitrant lipid A component [81,82]. On cell death, released LPS acts as a binding agent between soil particles, enhancing soil stability and water retention [83]. These adaptations highlight how microbial communities persist and exploit resources under carbon deficiency while shaping key soil ecosystem processes like organic carbon cycling and storage (Figs. 1 and 2).

While we focus on bacterial and archaeal regulation of SOM cycling, our study also highlights the need for investigating abiotic processes and interactions with fungi and viruses, which are harder to disentangle using current metagenomic methods [18–21]. Assembling and annotating fungal metagenomes presents clear challenges, as evidenced by the smaller fraction of reads and the relatively low number of assembled contigs compared to those of bacterial or archaeal origin; however, the annotation of these contigs revealed the potential role of soil fungal communities in contributing to soil carbon flux. At the same time, the multi-omics integration strategies employed in this study have facilitated the recovery of bacterial and archaeal lineages with unique yet complementary carbon sequestration strategies. However, FBA-derived metabolic inferences assume microbes operate in a steady state, and the incomplete recovery of MAGs may limit interpretation, warranting validation of these inferred traits using transcriptomics and/or proteomics; in particular, validation through cultivation methods and emerging computational tools for co-assemblies [84,85] would enhance the robustness of the findings. The recovery of 828 MAGs—though indicative of substantial prokaryotic diversity—may be biased toward ubiquitous, abundant taxa favored by continental-scale sampling across geographic gradients, and site-specific specialists and rare taxa involved in specialized carbon transformations [86,87] are likely underrepresented, suggesting that the results primarily reflect the “core” microbial carbon cycling machinery rather than the full functional diversity present in soil communities.

Nonetheless, our study provides a comprehensive, genome-resolved view of microbial decomposition potential across diverse soil environments, illuminating the metabolic underpinnings of carbon turnover at continental scale. By integrating thousands of chemically distinct SOM molecules with hundreds of metagenome-assembled genomes, we uncover widespread microbial capacities to degrade carbon compounds long thought to be chemically-resistant to decay. This work helps fill a critical knowledge gap in linking microbial taxonomy and metabolism to specific carbon pools, offering a mechanistic foundation for improving predictions of soil carbon dynamics under shifting environmental conditions. As global change accelerates, incorporating these microbial processes into Earth system models will be essential to more accurately project the fate of soil carbon reservoirs.

## Methods

### Data Collection

Through the 1000 Soils Pilot of the Molecular Observation Network (MONet) [14], 109 soil samples were collected from 60 sites across the CONUS. Full methodological details are available in refs. ([14] and [88]), and at https://github.com/EMSL-MONet/MONet-Protocols-. With a goal to investigate the microbial pathways that underscore variation in microbial activity across geography and soil depth profiles, we selected the subset of 47 soil samples, which exhibited the highest and lowest (30%) rates of respiration within either surface or subsoils for analysis. These soil samples spanned 37 geographical locations (Table 1) and were divided into four groups based on depth and respiration: surface-high (n=14), surface-low (n=12), suboil-high (n=11), and subsoil-low (n=10).

### Fourier Transform Ion Cyclotron Resonance Mass Spectrometry (FTICR-MS)

Water-soluble SOM composition was analyzed by Fourier Transform Ion Cyclotron Resonance Mass Spectrometry (FTICR-MS) [88,89]. Briefly, 6 g of dried soil in triplicate were resuspended in 30 mL DI water, and shaken for 2 hours at 800 rpm. The samples were then centrifuged at 6,000 rpm for 8 minutes and 5 mL of supernatant was transferred for solid phase extraction (SPE) with Agilent Bond Elut PPL cartridges [89] on Gilson ASPEC® SPE system. We used a Bruker 7 Tesla scimaX FTICR-MS at the Environmental Molecular Sciences Laboratory (EMSL) in Richland, WA to analyze the extracted SOM, with a negative ionization mode and ion accumulation time of 0.01 or 0.025 seconds. The mass accuracy was below 1 ppm, as confirmed by the Suwannee River Fulvic Acid (SRFA) samples. We used CoreMS (v. 2.0.0) [90] to process the raw FTICR-MS data, with noise thresholding of 10 (*noise_threshold_log_nsigma*) and minimum peak prominence of 0.1 (*peak_min_prominence_percent*). Mass calibration was performed by setting a threshold that uses the most calibration points (usually >200). Molecular formulae were annotated by both accurate mass and isotopologues, with a confidence score calculated for each formula. We then filtered the assigned peaks by *m/z* between 200 to 1,000, present in at least 2 out of 3 replicates, not present in two or more lab blanks, and with formulae confidence scores (combines *m/z* error and isotopic pattern) above 0.5 [91].

### Distribution and uniqueness of SOMs across the soil samples

Using the customized Python script, we assigned the 66,727 FTICR-derived molecular formulas into SOM compound classes, based on thresholds of O:C and H:C ratios on the van Krevelen diagram [92]. The matrix of the SOM assignment and the presence/absence matrix of the molecules across the 47 samples were used in a statistical analysis using the R package ftmsRanalysis [93] and associated scripts, to first determine the distribution of SOM types across the four soil sample types, and then to conduct a series of pairwise G-tests of uniqueness [94] with unadjusted p-value threshold of 0.05, generating profiles of SOMs that are differentially present in one soil type versus the other. The results of the G-tests were plotted as shown in Fig. 2.

### DNA extraction and sequencing

DNA was extracted using *Quick*-DNA Fecal/Soil Microbe Miniprep Kit D6010 (ZymoResearch) and then cleaned and concentrated with DNA Clean and Concentrator-25 (ZymoResearch). Samples were sequenced at Azenta Life Sciences using the Illumina Whole Genome Metagenomics pipeline or at the Joint Genome Institute, following identical protocols.

### Quality control, assembly and binning of the metagenome reads

Metagenome construction, including quality-control/read filtering, contig assembly and binning processes, were performed using a suite of tools available in the MetaGEM pipeline [95]. Specifically, short read quality filtering and adapter trimming were applied to the set of reads (de-interleaved) using fastp v0.20.0 [96] with default settings. The filtered reads from each sample were assembled into contigs using MEGAHIT v1.2.9 [97], with *--presets meta-sensitive* and *--min-contig-len 1000* flags.

Before the binning process, contig coverages across the samples (required by some of the binning tools as described below) were generated by cross-mapping the filtered short reads to the set of assemblies. This was accomplished by generating an index for each contig using the ‘bwa index’ command with default settings. The set of indices were used as part of the input for bwa-mem v0.7.17 [98], generating SAM files, which were subsequently converted to BAM format using the ‘samtools view’ command with the *-Sb* flag, and then were sorted using the ‘samtools sort’ command with the default settings. The sorted BAM files were then used to generate the contig coverages for the binning tools, MetaBAT2 v.2.12.1 [99] and MaxBin2 v2.2.5 [100], using the script, ‘jgi_summarize_bam_contig_depths’ available as part of the MetaBAT2 package. The contig coverage files were then used as input data for the MetaBAT2 and MaxBin2, with default settings for each tool.

Binning was performed using the tool CONCOCT v1.1.0 [101] and the scripts available in the MetaGEM pipeline. The coverage files were generated using the script, ‘concoct_coverage_table.py’ (default settings), with a set of BED files generated for each contig as input. The BED files were created using the ‘cut_up_fasta.py’ script, with the parameters *-c 10000 -o 0 -m b* that split the contigs into 10 kb fragments. For each fragment, CONCOCT was run using the coverage file as input and with the *-c 800* parameter, yielding a set of bins for each sample, based on the split contigs. The script ‘extract_fasta_bins.py’ was then run to generate bins using the original unfragmented contigs.

### Refinement and reassembly of the MAGs

To extract the best version of bins from the outputs of the three binning tools, the ‘bin_refinement’ script available in metaWRAP v1.2.3 [102] was used with *-x 10 -c 50* parameters. This command de-replicates the results of the three binning methods by selecting the bin with the highest CheckM2 [103] completeness and the lowest contamination, with the completeness taking precedence over contamination. The parameters in this script ensure that only the binning results with contamination and completeness thresholds of ≤ 10% and ≥ 50%, respectively, are considered for the bin refinement process.

To improve the quality of the bins selected from the refinement step, the ‘reassemble_bins’ command (with the same parameters used for the ‘bin_refinement’ script) was used. This script extracts the reads that map to the contigs belonging to the refined bins and reassembles them using the SPAdes assembly. This process is done at both a ‘strict’ (no mismatches) or ‘permissive’ (< 5 mismatches) level. CheckM2 completeness and contamination between the three versions of bins (i.e., strict, permissive or original derived from the refinement) are compared, and the bins with the best score are selected as the finalized MAGs, yielding 344 genomes.

To determine the distribution of the bacterial and archaeal communities that were represented by the MAGs, we first used the ‘pipe’ subcommand in SingleM [104], generating a summary of operational taxonomic units (OTUs) detected based on 35 bacterial and 37 single-copy marker genes identified from MAGs. The OTU summary table was used as input for the ‘appraise’ function in SingleM, employing the parameters *--imperfect* and *--sequence-identity 0.87* to account for genomes that are similar to those found in the metagenome that were classified at the genus level (i.e. average nucleotide identity of 87%) [105].

According to the SingleM appraise result, the 344 MAGs represented only 44.4% and 32.3% of the bacterial and archaea diversity. Thus, we made an attempt to improve the MAG recovery through coassembly of the reads, as implemented in Bin Chicken [84]. Specifically, we used this tool to identify MAGs through co-assembly of the reads from the soils associated with each of the four depth- and respiration-level classification. We first executed Bin Chicken with the single subcommand to generate MAGs from each sample, using forward and reverse reads and SingleM-generated files (e.g., protein-coding genes and two OTU tables—one from contigs and one from MAGs) as input, with the parameters *--run-aviary* and *--assembly-strategy* megahit. The output MAGs from the ‘single’ subcommand were used to execute two rounds of iterative coassemblies via the ‘iterate’ subcommand. For each run of the ‘iterate’ function, we used the same set of parameters as we did with the ‘single’ subcommand. By the end of the second execution of ‘iterate,’ the total number of MAGs increased to 828, improving the SingleM appraisal value to 50.6% and 36.5% for bacteria and archaea. Dereplication of the MAGs and calculating genome abundances were performed using the methods described below.

### Genome dereplication

The 828 bins that were generated were dereplicated using galah (v0.4.0) [106] at 95% average nucleotide identity (ANI) similarity, which can be used to represent species-level similarities [107], resulting in 358 clusters. Representative, dereplicated MAGs from each cluster were selected based on the quality metric derived from CheckM2.

### Genome abundances

The abundance of the dereplicated MAGs in each of the samples was calculated via mapping of the filtered reads to the dereplicated MAG dataset using CoverM [108] with the following flags: *-p minimap2-sr -m trimmed_mean --min-read-percent-identity 0.80 --min-read-aligned-percent 0.75 --min-covered-fraction 20*. Read mapping indicated that the MAGs accounted for 1.3% to 22.2% of the reads of each sample.

### Phylogenetic assignments of dereplicated MAGs

The dereplicated MAGs generated using the methods described above were taxonomically classified using the ‘identify’, ‘align’ and ‘classify’ commands available in GTDB-Tk v2.5.2 [109]. This set of commands identifies 120 ubiquitous bacterial marker genes (and 53 archaeal genes) from the input MAGs, which are then concatenated and aligned together. The alignment of the concatenated marker genes is then used as input for Pplacer [110] to assign each MAG into the optimal loci in the bacterial or archaeal tree of life. The output of the process includes tabular files containing taxonomic classifications of the MAGs. For each of the depth and respiration-level classifications, the average abundances of the taxonomic groups were calculated from the total abundances of the dereplicated MAGs assigned to each lineage.

### Assembly and analysis of fungal metagenomes

We used Kraken2 [22] to recruit the DNA reads against the fungal taxonomic marker genes present in the RefSeq Fungi [111] and the 18S SSU rRNA sequences from the SILVA database [112]. On average, 1.33x10^5^ fungal reads were recovered, making up < 1% (i.e., 0.24%) of the total reads from each soil. The reads were assembled using MEGAHIT v1.2.9 with parameters *--presets meta-sensitive --min-contig-len 1000*, generating an average of 86 contigs per sample. The presence of the carbohydrate-active enzymes (CAZymes) in the fungal contigs was assessed by searching them against the Carbohydrate Active Enzymes (CAZy) database [29], using dbCAN2 [113].

### KEGG pathway analysis of FTICR-derived molecules using KO identifications

The ftmsRanalysis package and associated custom scripts mentioned above were also used to generate a summary of KO identifications that were mapped to the FTICR-derived molecules. These summary tables were parsed to extract the KOs present in 50% or more of the samples belonging to each partition. The KEGG pathway representation analysis, which employs the enrichKEGG algorithm in the clusterProfiler R package [114], was conducted using this set of KOs. This method calculates the ratio of KOs affiliated with specific pathways versus the total number of KOs in each sample set. This analysis generated a summary visualization of significant KEGG pathways present in each layer/respiration partition (Fig. 4).

### Network analysis of the relationship between KO derived from functional annotation of MAGs and FTICR-MS derived molecules

For each of the four layer/respiration partitions, co-occurrence networks were constructed using Spearman’s correlation to explore the relationship between the KOs detected from the MAGs and the FTICR-MS-derived compounds. To generate the KO abundance profile, functional genes were first detected from the nucleotide sequences of all dereplicated MAGs using Prokka [115], and were searched against KEGG Orthology (KO) [116] using the ‘annotate’ function implemented in EnrichM [117]. For the bacterial MAGs, Prokka was executed with *--kingdom Bacteria* and *--addgenes* parameters, while for the archaeal MAGs, the *--kingdom Archaea* flag was used instead. The abundance tables of KO and the compounds were used as input for Spearman correlation implemented in R. Each correlation matrix was filtered to obtain relationships with *rho* > 0.6 and FDR-corrected P < 0.01 [118]. The output matrices were then imported into Cytoscape [119] version 3.10.3. Clusters in each partition were identified using the MCODE application, with default parameters, producing a set of clusters with scores representing their densities and interconnectedness [120].

### Assessment of microbial carbohydrate depolymerization in abundant prokaryotes using CAZy annotation of MAGs

We selected two order-level bacterial lineages, *Rhizobiales* and *Chthoniobacterales*, each of which is represented by > 5.0% of the 319 bacterial MAGs recovered from all soil samples. Only one order-level archaeal lineage (i.e., *Nitrososphaerales*) was selected, as > 90% of the 39 MAGs were assigned to this group. Functional genes detected from MAGs of these bacterial and archaeal clades using Prokka (see above) were searched against the Carbohydrate-Active Enzymes (CAZy) database [29], using dbCAN2 [113] to predict the capacities of these lineages to depolymerize lignin and other chemically recalcitrant carbohydrates.

### Metabolic model prediction for Rhizobiales, Chthoniobacterales and Nitrososphaerales

Using the functional annotation profiles generated from the method described in the previous section, we inferred metabolic model predictions for the two dominant bacterial orders, *Rhizobiales* and *Chthoniobacterales* and the archaeal order *Nitrosophaerales*. To accomplish this, we built a metabolic prediction narrative using a selected suite of applications available in the DOE Systems Biology Knowledgebase (KBase) [121] platform. In this narrative, GFF tables generated using Prokka containing genomic feature information (i.e., gene names, enzyme commission identifications and protein names; see above section) for each detected coding region and the nucleotide FASTA files of the MAGs are used to re-annotate the genomes using RAST-Tk v1.73 (Rapid Annotation using Subsystems Technology - Toolkit) [122]. This process generates the output files compatible with the KBase metabolic modeling applications.

The RAST-Tk annotation objects were used as input for the ModelSEED [123] application, generating models representing the broader metabolic properties in the bacterial and archaeal lineages under investigation. The models were generated by compiling the i metabolic pathways information encoded by genes detected in the MAGs. The output generated by ModelSEED includes a matrix of metabolic reactions, their associated biochemical compounds, and the gene-protein-reaction (GPR) that encompass the dependencies between the responses and the related genes. The GPR can help differentiate between cases where a reaction is catalyzed by a protein complex encoded by multiple genes and where several individual proteins independently orchestrate a reaction. The ModelSEED output also includes a ‘biomass objective function’ file that indicates the abundance of the metabolites generated by the microbes (e.g., amino acids, lipids and quinones) as bi-products of their growth metabolism.

Finally, the Flux Balance Analysis (FBA) [124] application was used to read the ModelSEED-derived metabolic models and a default media file provided by KBase that specified the set of > 500 compounds an organism can use for its growth, generating tables of metrics that allowed estimation of growth rates of microbes and/or output of specific metabolites on a specified media. The FBA estimates the growth rate of an organism by calculating growth-optimal fluxes through all reactions in the metabolic network. The uptake values for the compounds specified in the output file, measured in mmol per gram cell dry weight per hour, indicate whether the microbes use the metabolites for growth or excrete them as a by-product. The list of the output uptake values was examined to determine the spectrum of compounds likely to be metabolized by the members of the prevalent order-level clades detected in the soils.

## Data availability

The sequencing data associated with this study are available in the NCBI Sequence Read Archive (SRA) under accession number **PRJNA1260013**. Data in the MONet open science database can be found at https://sc-data.emsl.pnnl.gov/monet and MONet Zenodo DOI (https://doi.org/10.5281/zenodo.7406532).

## Code availability

The R and Python code used for analyzing the FTICR-MS data and for preparing the main and supplementary figures are available upon request at the following GitHub repository: https://github.com/EMSL-MONet/The-1000-Soil-ICR-Metagenome. The KBase narrative developed to analyze the *Rhizobiales* and *Chthoniobacterales* MAGs can be accessed at https://narrative.kbase.us/narrative/186750.

## Acknowledgements

This research was performed on a project award DOI https://doi.org/10.46936/intm.proj.2021.60141/60000423 from the Environmental Molecular Sciences Laboratory (EMSL), a DOE Office of Science user facility sponsored by the Biological and Environmental Research program under Contract No. DE-AC05-76RL01830. Soil data was provided by the Molecular Observation Network (MONet) at the EMSL (DOI: https://doi.org/10.46936/10.25585/60008970). MONet is conducted in collaboration with Joint Genome Institute (https://ror.org/04xm1d337), a DOE Office of Science user facility supported by the Office of Science of the U.S. Department of Energy operated under Contract DE-AC02-05CH11231. The MONet database is an open, FAIR, and publicly available compilation of the molecular and microstructural properties of soil. The National Ecological Observatory Network is a program sponsored by the U.S. National Science Foundation and operated under cooperative agreement by Battelle. A subset of samples used in this research were obtained through NEON Research Support Services.

---

**Supplementary Figure 1.**
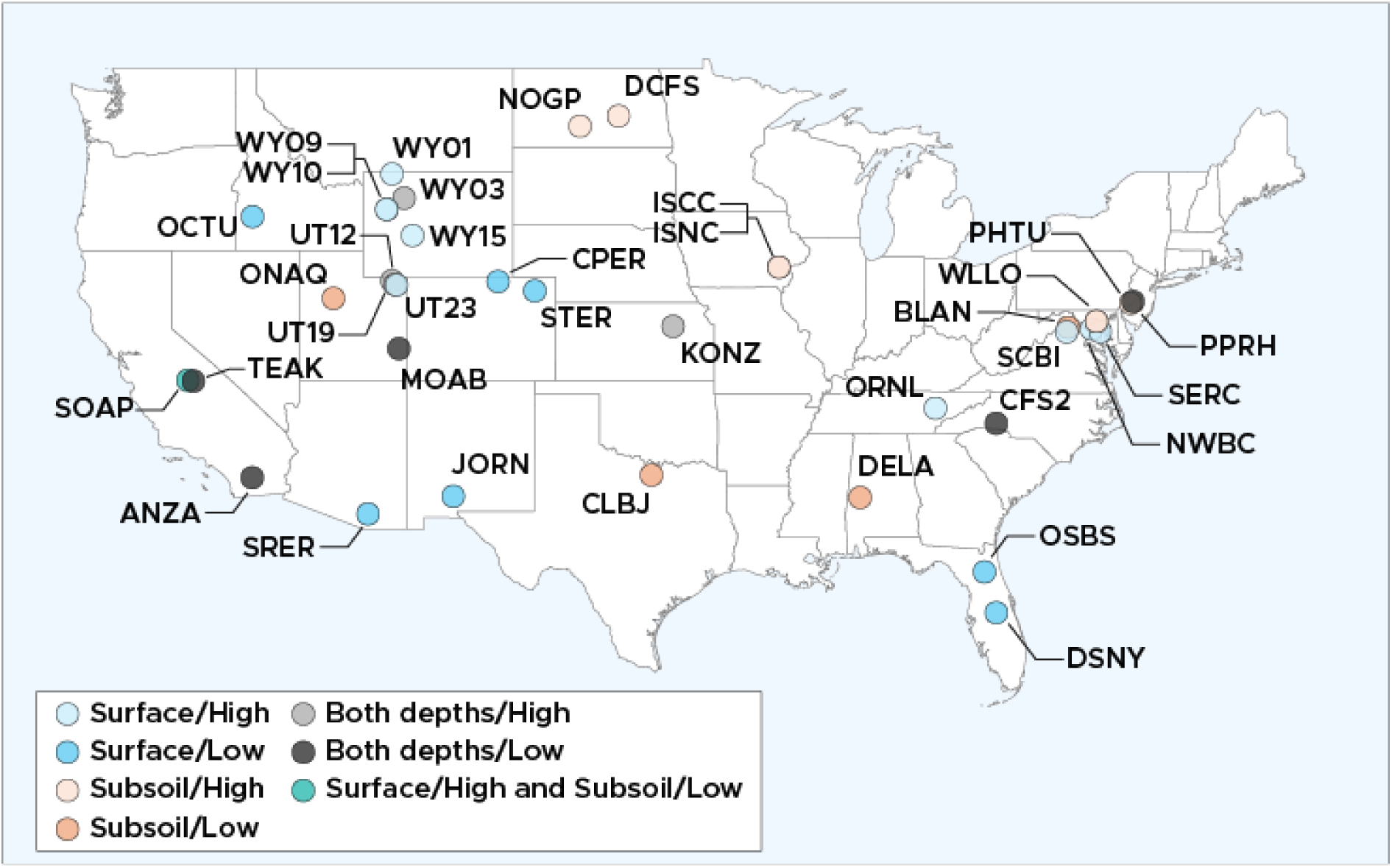
Geographic coordinates of 47 soil samples spanning 37 sites across the continental United States (CONUS). Sample types, defined by both soil depths and respiration levels, are represented by color codes as specified in the legend. The map and geographic coordinates were generated using the rnaturalearth, ggplot2, and dplyr R packages.

**Supplementary Figure 2.**
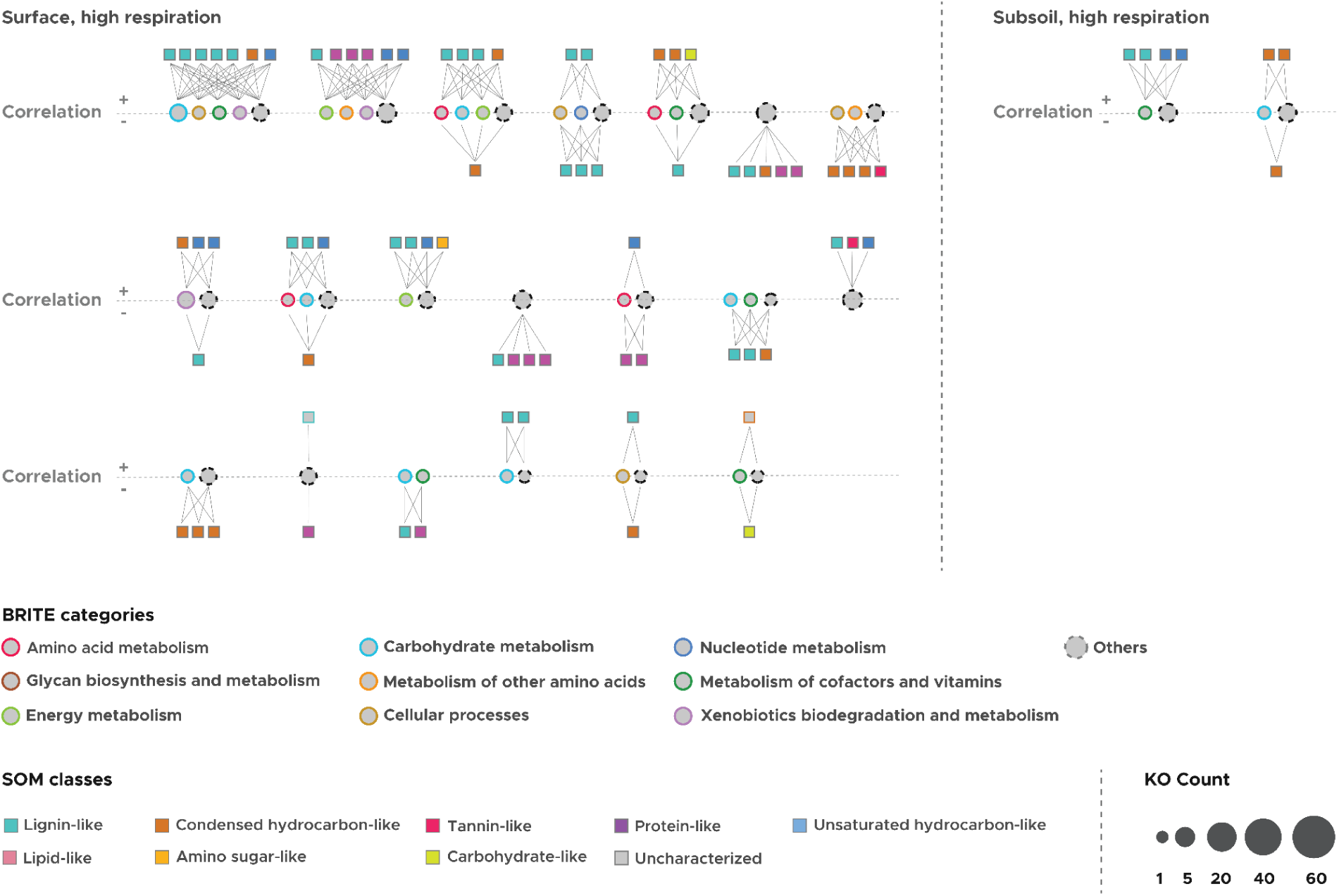
Modules with significant correlations between SOMs and KEGG Orthology (KO)-annotated genes detected in high-respiration soils, with the MCODE cluster score of < 5. The shapes of the nodes within each module signify either KOs or SOMs, with their colors representing KEGG pathway categories for KOs or SOM classes, as detailed in the legend. Positive correlations between SOMs and KOs are depicted as edges situated above the dashed “Correlation” line, whereas edges indicating negative correlations are positioned below the dashed line. These correlations meet the criteria of rho > 0.6 and FDR-corrected P < 0.01.

**Supplementary Figure 3.**
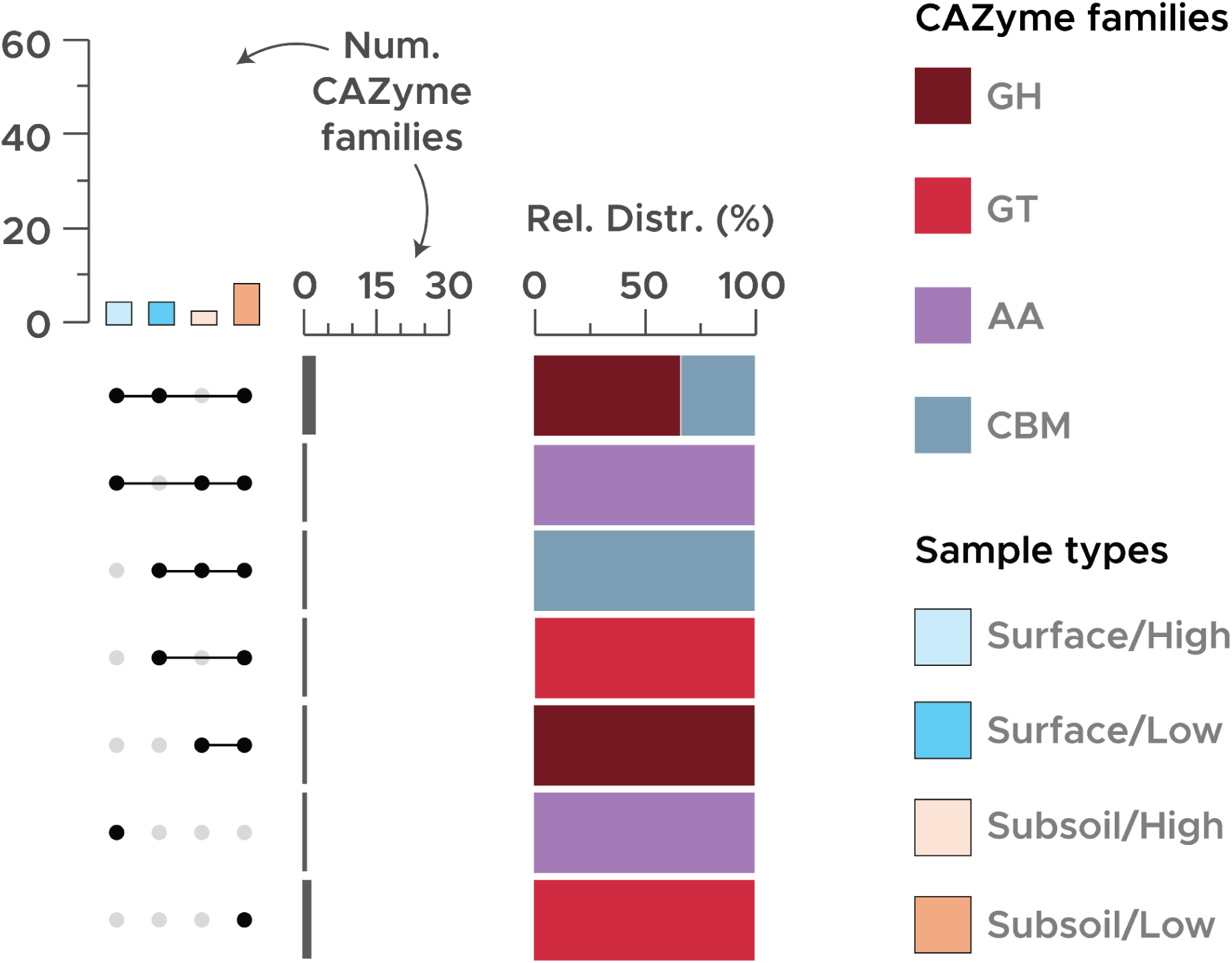
Carbohydrate-active enzymes (CAZymes) detected in fungal contigs. The bar chart on the top illustrates the total number of CAZyme families identified from the contigs derived from the surface or subsoils under high or low respiration. Meanwhile, the combined dot-and-line plot along with the bar chart on the left displays the count of CAZyme families found in one or more of the soil sample types. Additionally, the stacked bar chart highlights the proportional distribution of CAZyme categories among these families. The CAZyme categories, as defined in the legend, include GH (glycoside hydrolase), GT (glycosyl transferase), CE (carbohydrate esterase), AA (auxiliary activity), CBM (carbohydrate-binding module), and PL (polysaccharide lyase).

