## Supplementary material for "Continental-scale integration of soil metagenomes and organic matter chemistry reveals ubiquitous microbial capacity for chemically-recalcitrant carbon decomposition": Table 1

**Table 1**. Distribution of CAZyme classes detected in selected genera affiliated with the order-level lineages, *Rhizobiales*, *Chthoniobacterales* and *Nitrososphaerales*. The percentages next to the counts of each class represent the relative distribution of total classes for each lineage, as indicated by the numbers in parentheses beside their names.

|  | *Rhizobiales* | | | | *Chthoniobacterales* |
| --- | --- | --- | --- | --- | --- |
| CAZyme class | *Methyloceanibacter* (43) | *Bradyrhizobium* (60) | *Pseudolabrys* (52) | VAZQ01 (54) | *Udaeobacter* (28) |
| GH | 22 (51.2%) | 30 (50.0%) | 30 (57.7%) | 25 (46.3%) | 12 (42.8%) |
| GT | 10 (23.2%) | 17 (28.3%) | 9 (17.3%) | 17 (31.5%) | 8 (28.6%) |
| AA | 2 (4.6%) | 5 (8.3%) | 6 (11.5%) | 4 (7.4%) | 4 (14.3%) |
| CBM | 4 (9.3% | 2 (3.3%) | 2 (3.8%) | 3 (5.6%) | 1 (3.6%) |
| CE | 5 (11.6%) | 6 (10.0%) | 5 (9.6%) | 5 (9.2%) | 3 (10.7%) |
| PL | 0 (0.0%) | 0 (0.0%) | 0 (0.0%) | 0 (0.0%) | 0 (0.0%) |

|  | *Nitrososphaerales* | | |
| --- | --- | --- | --- |
| CAZyme class | JAFAQB01 (52) | *Nitrososphaera* (92) | TA-21 (53) |
| GH | 28 (53.8%) | 49 (53.3%) | 22 (41.5%) |
| GT | 16 (30.8%) | 21 (22.8%) | 17 (32.1%) |
| AA | 1 (1.9%) | 5 (5.4%) | 5 (9.4%) |
| CBM | 3 (5.8%) | 8 (8.7%) | 2 (3.8%) |
| CE | 4 (7.7%) | 7 (7.6%) | 6 (11.3%) |
| PL | 0 (0.0%) | 2 (2.2%) | 1 (1.9%) |
